# The I*n* S*ilico* Genotyper (ISG): an open-source pipeline to rapidly identify and annotate nucleotide variants for comparative genomics applications

**DOI:** 10.1101/015578

**Authors:** Jason W. Sahl, Stephen M. Beckstrom-Sternberg, James S. Babic-Sternberg, John D. Gillece, Crystal M. Hepp, Raymond K. Auerbach, Waibhav Tembe, David M. Wagner, Paul S. Keim, Talima Pearson

## Abstract

The identification and annotation of nucleotide variants, including insertions/deletions and single nucleotide polymorphisms (SNPs), from whole genome sequence data is important for studies of bacterial evolution, comparative genomics, and phylogeography. The *in* S*ilico* Genotyper (ISG) represents a parallel, tested, open source tool that can perform these functions and scales well to thousands of bacterial genomes. ISG is written in Java and requires MUMmer (Delcher, et al., 2003), BWA (Li and Durbin, 2009), and GATK (McKenna, et al., 2010) for full functionality. The source code and compiled binaries are freely available from https://github.com/TGenNorth/ISGPipeline under a GNU General Public License. Benchmark comparisons demonstrate that ISG is faster and more flexible than comparable tools.

## INTRODUCTION

The application of next generation sequencing (NGS) technologies to microbiology has changed our view of bacterial evolution and relatedness. Large-scale phylogeographic studies have been conducted for bacterial species including *Bacillus anthracis* (Keim and Wagner, 2009). The correlation of geographic distribution with NGS data from bacterial isolates has largely focused on the analysis of single nucleotide polymorphisms (SNPs). Although many groups have used SNP discovery to generate phylogenetic trees (Pandya, et al., 2009), the methods that have been employed remain largely unpublished, tested, and validated. Here we present the *in silico* genotyper (ISG), an open-source tool that can be used for SNP and inversion/deletion (indel) discovery, annotation, and phylogenomics.

## IMPLEMENTATION

ISG is written in Java and relies on the Queue pipeline system from the Broad Institute (http://www.broadinstitute.org/gatk/auth?package=Queue) for pipeline execution. ISG can handle multiple types of input data, including raw reads in either “.txt”, “.fastq”, or “.fastq.gz” format, genome assemblies in “.fasta” format, genome annotation in “.gbk” format, binary alignment map (“.bam”) files, and/or variant call format (“.vcf”) files (Figure 1). If single or paired reads are provided, BWA-MEM (Li, 2013) can be used to align the reads against a reference genome in “.fasta” format (the reference genome can either be a finished genome or a locally generated draft genome assembly). SNPs and indels are called with the UnifiedGenotyper method in GATK (McKenna, et al., 2010), using user-defined thresholds for minimum depth of coverage and allele proportion variation. All variables used by GATK can be modified by the user for specific applications. If external genome assemblies are supplied, ISG calls SNPs using the show-snps function in MUMmer (Delcher, et al., 2002). When a SNP is called from the raw reads or assembly in at least one genome, that position is queried in all other genomes. If a position in a query genome fails to pass one of the user-defined filters (e.g. minimum coverage depth), an “N” is applied to that position. If a position is not a SNP in a query genome and it passes all filters, an additional test is conducted by GATK (callableLoci) to quantify the base quality and coverage at that position; a position of sufficient quality, as determined by GATK, is assumed to be the reference state. Otherwise, a ‘.’ or ‘N’ is substituted at that position depending on sufficient coverage or quality, respectively.

**Figure 1.**
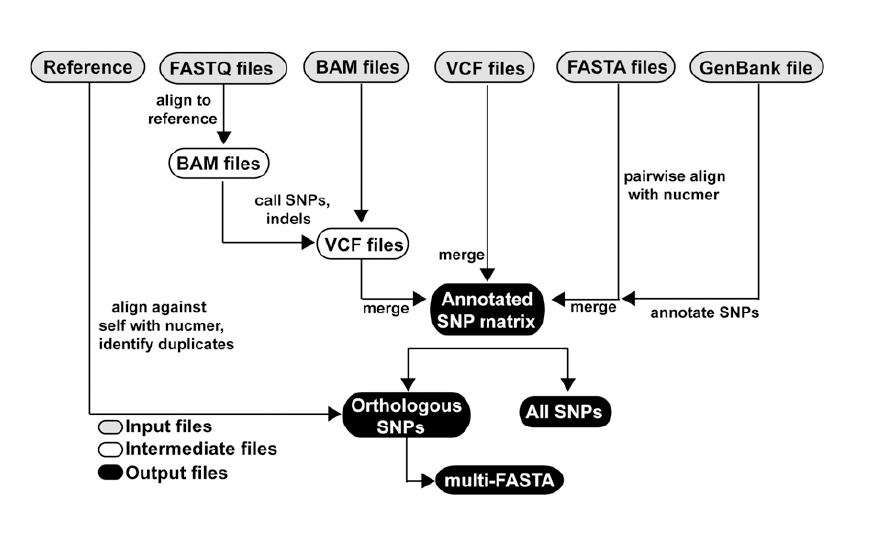
A workflow of the ISG pipeline

## COMPARISON WITH OTHER METHODS

Other software tools have been published in order to use SNP data to infer phylogenies and annotate SNPs. The snpTree method (Leekitcharoenphon, et al., 2012) was published and performs similar functions to ISG. However, snpTree is a web-based server, and while useful for a small number of genomes, it is currently impractical for uploading raw sequence data for hundreds to thousands of bacterial genomes. We attempted to test the results of snpTree, but we could not get even moderate sized datasets to complete successfully. SPANDx (Sarovich and Price, 2014) is a reference dependent approach that can also produce SNP matrices from raw sequence reads. However, SPANDx cannot process genome assemblies and is reliant on the Torque queuing system, which limits its flexibility, and was therefore not compared with ISG. Other, reference independent approaches, including kSNP 2 (Gardner and Hall, 2013) and CO-Phylog (Yi and Jin, 2013), have also been published. CO-Phylog produces a distance matrix as output, but does not perform SNP annotation; this output limits the types of phylogenetic analyses that can be performed. kSNP can process either assembled genomes or raw reads, although the raw reads need to be concatenated, which increases the initial processing time. The functionality of kSNP is similar to ISG in terms of SNP identification and annotation, however the method for the identification of SNPs between methods is very different. ParSNP is a method that can rapidly compare the core genome between a large number of genome assemblies (Treangen, et al., 2014); however, ParSNP can currently not process raw data and requires an additional assembly step.

To test the speed of the different methods, 1000 *Escherichia coli* and *Shigella* genomes were downloaded from Genbank and randomly subsampled at 100 genome intervals, from 100 to 900 genomes; the subsampled genomes were then processed with kSNP, CO-Phylog, and ISG, all using 16 threads, when available. Speed tests demonstrate that ISG scales linearly with an increasing number of genomes (Figure 2). kSNP failed at 400 genomes and the CO-Phylog sampling was stopped at 300 genomes due to prohibitive time requirements.

**Figure 2.**
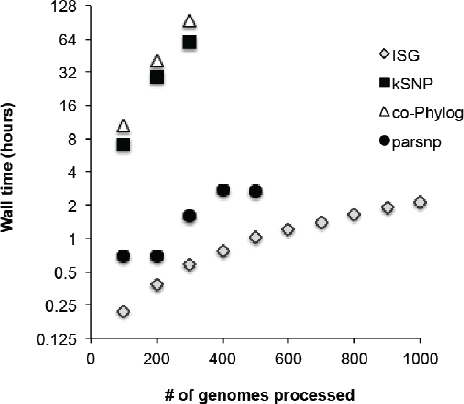
A time comparison of ISG with other methods. Each method was run with 16 processors, when available.

To test the genotyping reproducibility of the ISG algorithm, both raw reads and assemblies from 118 *Yersinia pesti*s genomes (Cui, et al., 2012) were downloaded from Genbank (Supplemental Table 1); an additional 15 genome assemblies were also downloaded (Supplemental Table 1). A recent paper analyzed assemblies from these genomes and characterized 2,298 non-homoplastic SNPs. Although the method that identified these SNPs could not be reproduced, ISG was run on these assemblies to determine how many of these SNPs could be identified. Using the assemblies submitted to GenBank (n=133), 2,078 of the SNPs were called by ISG and 2,233 were called by kSNP. From using a combination of genome assemblies (n=15) and raw reads (n=118), 1,939 of the SNPs were called by ISG, while 2,037 were called by kSNP. Although this discrepancy between using raw reads or assemblies is likely due to the presence of assembly errors, without the source code for the other method, a true comparison cannot be performed. SNPs only present in the Cui *et al*. analysis were manually checked in the short read alignments and were found to all be monomorphic. ISG called an additional 9 SNPs that were manually confirmed to be present in this dataset, but were missing from the Cui *et al*. analysis (Table 1); two of these 9 SNPs were identified by kSNP from an analysis of assemblies alone while 7 of these 9 SNPs were identified by ISG from assemblies alone.

**Table 1.** Details of SNPs identified with different methods and data types

| Chromosome | position | Reference state | SNP in Cui et al.? | SNP with kSNP (assemblies)? | SNP with ISG (reads+assemblies) | SNP with ISG (assemblies) |
| --- | --- | --- | --- | --- | --- | --- |
| NC_003143.1 | 229742 | C | no | yes | yes | yes |
| NC_003143.1 | 4491401 | T | no | no | yes | yes |
| NC_003143.1 | 4556781 | T | no | no | yes | no |

## CONCLUSION

SNP and indel discovery and annotation are important data required for comparative genomics, phylogeography, and functional studies. Tools that can accurately and rapidly perform these functions are required, especially as the number of sequenced genomes rapidly increases. ISG represents an open source, parallel, tested method for whole genome comparative genomics and phylogenomics.

## ACKNOWLEDGEMENTS

John Pearson, Jicheng Hao (deceased), Shripad Sinari, assisted with development of the algorithms. Jeffrey Foster, Rich Okinaka, Dawn Birdsell, Heidie Hornstra O’Neill, Amy Vogler, Erin Price, Derek Sarovich, Kevin Drees, James Schupp, Elizabeth Driebe, Lance Price, supplied data and feedback for results.

*Funding*: TGen, US Army Medical Research Contract W81XWH-07-1-0100 and NIH grant 1R15AI075333-01 to SBS. NIH 1 UO1 AI066581-01 to PK. US Department of Homeland Security Science and Technology Directorate (award HSHQDC-10-C-US).

## References

Cui, Y., et al. Historical variations in mutation rate in an epidemic pathogen, *Yersinia pestis*. Proc Natl Acad Sci U S A 2012;110(2):577-582.

Delcher, A.L., et al. Fast algorithms for large-scale genome alignment and comparison. Nucleic Acids Res 2002;30(11):2478-2483.

Delcher, A.L., Salzberg, S.L. and Phillippy, A.M. Using MUMmer to identify similar regions in large sequence sets. Curr Protoc Bioinformatics 2003;Chapter 10:Unit 10 13.

Gardner, S.N. and Hall, B.G. When whole-genome alignments just won’t work: kSNP v2 software for alignment-free SNP discovery and phylogenetics of hundreds of microbial genomes. PLoS ONE 2013;10.1371/journal.pone.0081760.

Keim, P.S. and Wagner, D.M. Humans and evolutionary and ecological forces shaped the phylogeography of recently emerged diseases. Nature reviews. Microbiology 2009;7(11):813-821.

Leekitcharoenphon, P., et al. snpTree - a web-server to identiy and construct SNP trees from whole genome sequence data. BMC genomics 2012;13((Suppl 7):S6).

Li, H. Aligning sequence reads, clone sequences and assembly contigs with BWA-MEM. arXiv.org 2013(1303.3997 [q-bio.GN]).

Li, H. and Durbin, R. Fast and accurate short read alignment with Burrows-Wheeler transform. Bioinformatics 2009;25(14):1754-1760.

McKenna, A., et al. The Genome Analysis Toolkit: a MapReduce framework for analyzing next-generation DNA sequencing data. Genome Res 2010;20(9):1297-1303.

Pandya, G.A., et al. Whole genome single nucleotide polymorphism based phylogeny of Francisella tularensis and its application to the development of a strain typing assay. BMC microbiology 2009;9:213.

Sarovich, D.S. and Price, E.P. SPANDx: a genomics pipeline for comparative analysis of large haploid whole genome re-sequencing datasets. BMC research notes 2014;7(1):618.

Treangen, T.J., et al. Rapid core-genome alignment and visualization for thousands of microbial genomes. bioRxiv 2014.

Yi, H. and Jin, L. Co-phylog: an assembly-free phylogenomic approach for closely related organisms. Nucleic Acids Res 2013;41(7):e75.

